# SomaticSignatures: Inferring Mutational Signatures from Single Nucleotide Variants

**DOI:** 10.1101/010686

**Authors:** Julian S. Gehring, Bernd Fischer, Michael Lawrence, Wolfgang Huber

## Abstract

Mutational signatures are patterns in the occurrence of somatic single nucleotide variants (SNVs) that can reflect underlying mutational processes. The *SomaticSignatures* package provides flexible, interoperable, and easy-to-use tools that identify such signatures in cancer sequencing data. It facilitates large-scale, cross-dataset estimation of mutational signatures, implements existing methods for pattern decomposition, supports extension through user-defined methods and integrates with Bioconductor workflows.

The R package *SomaticSignatures* is available as part of the Bioconductor project (R Core Team, 2014; Gentleman *et al*., 2004). Its documentation provides additional details on the methodology and demonstrates applications to biological datasets.

## 1 Introduction

Mutational signatures link observed somatic single nucleotide variants to mutation generating processes (Alexandrov *et al*., 2013a). The identification of these signatures offers insights into the evolution, heterogeneity and developmental mechanisms of cancer (Fischer *et al*., 2013; Alexandrov *et al*., 2013b; Nik-Zainal *et al*., 2012). Existing softwares offer specialized functionality for this approach and have contributed to the characterization of signatures in multiple cancer types (Nik-Zainal *et al*., 2012; Fischer *et al*., 2013), while their reliance on custom data input and output formats limits integration into common workflows.

The *SomaticSignatures* package aims to encourage wider adoption of mutational signatures in tumor genome analysis by providing an accessible R implementation that supports multiple statistical approaches, scales to large datasets, and closely interacts with the data structures and tools of Bioconductor.

## 2 Approach

The probability of a somatic SNV to occur can depend on the sequence neighborhood, and a fruitful approach is to analyze SNV frequencies together with their immediate sequence context, the flanking 3*′* and 5*′* bases (Alexandrov *et al*., 2013b). As an example, the mutation of `A` to `G` in the sequence `TAC` defines the mutational motif `T[A>G]C`. The occurrence patterns of such motifs capture characteristics of mutational mechanisms, and the frequencies of the 96 possible motifs across all samples define the mutational spectrum. It is represented by the matrix *M*_*ij*_, with *i* enumerating the motifs and *j* the samples. The mutational spectrum can be interpreted by decomposing *M* into two matrices of smaller size (Nik-Zainal *et al*., 2012),

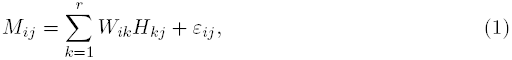

where the number of signatures *r* is typically small compared to the number of samples and the elements of the residual matrix *ε* are minimized, such that *W H* is a useful approximation of the data. The columns of *W* describe the composition of a signature: *W*_*ik*_ is the relative frequency of somatic motif *i* in the *k*-th signature. The rows of *H* indicate the contribution of each signature to a particular sample *j*. A primary goal of the *SomaticSignatures* package is the easy application of this approach to datasets in an environment that provides users with powerful visualisations and algorithms.

## 3 Methods

Several approaches exist for the decomposition (Eq. 1) that differ in their constraints and computational complexity. In principal component analysis (PCA), for a given *r*, *W* and *H* are chosen such that the norm 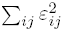 is minimal and *W* is orthonormal. Non-negative matrix factorization (NMF) (Brunet *et al*., 2004) is motivated by the fact that the mutational spectrum fulfills *M*_*ij*_ ≥ 0, and imposes the same requirement on the elements of *W* and *H*. Different NMF and PCA algorithms allow additional constraints on the results, such as sparsity. To deduce the number *r* of signatures present in the data, information theoretical criteria as well prior biological knowledge can be employed (Nik-Zainal *et al*., 2012; Alexandrov *et al*., 2013a).

## 4 Results

*SomaticSignatures* is a flexible and efficient tool for inferring characteristics of mutational mechanisms, based on the methodology developed by Nik-Zainal *et al*. (2012). It integrates with Bioconductor tools for processing and annotating genomic variants. An analysis starts with a set of SNV calls, typically imported from a VCF file and represented as a `VRanges` object (Obenchain *et al*., 2014). Since the original calls do not contain information about the sequence context, we construct the mutational motifs first, based on the reference genome.

`ctx = mutationContext(VRanges, ReferenceGenome)`

Subsequently, we define the mutational spectrum *M*. While its columns are by default constructed according to the sample labels, users can specify an alternative grouping covariate, for example drug response or tumor type.

`m = motifMatrix(ctx, group)`

Mutational signatures and their contribution to each sample’s mutational spectrum are estimated with a chosen decomposition method for a defined number of signatures. We provide convenient access to implementations for NMF and PCA (Gaujoux *et* Seoighe, 2010; Stacklies *et al*., 2007), and users can apply own functions with alternative decomposition methods through the API.

`sigs = identifySignatures(m, nSig, method)`

The user interface and library of plotting functions facilitate subsequent analysis and presentation of results (Fig. 1). Accounting for technical biases is often essential, particularly when analyzing across multiple datasets. For this purpose, we provide methods to normalize for the background distribution of sequence motifs, and demonstrate how to identify batch effects.

**Figure 1:**
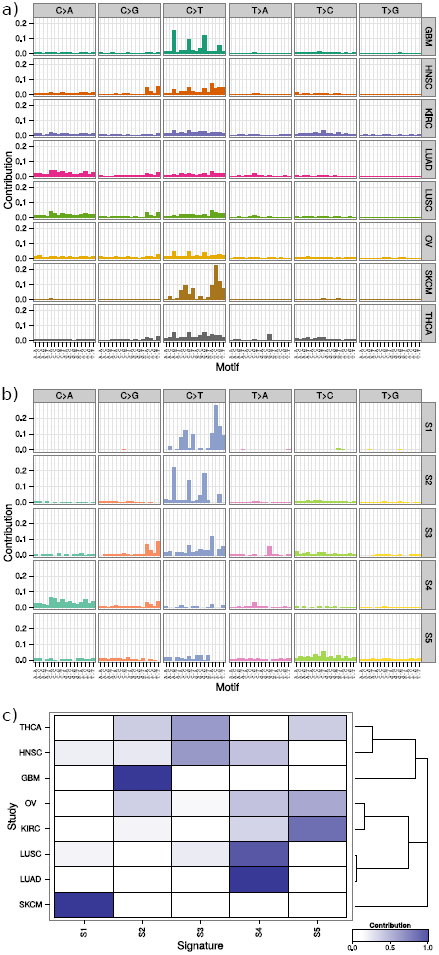
Analysis of mutational signatures for eight TCGA studies (Gehring, 2014). The observed mutational spectrum of each study (panel a) was decomposed into 5 distinct mutational signatures S1 to S5 (panel b) with NMF. The presence of these signatures in the studies (panel c), as shown by hierachical clusting, underlines the similarities in mutational processes of biologically related cancer types. An annotated high-resolution version of this figure is available as Supplementary Figure S1.

In the documentation of the software, we illustrate a use case by analyzing 594,607 somatic SNV calls from 2,408 TCGA whole-exome sequenced samples (Gehring, 2014). The analysis, including NMF, PCA and hierarchical clustering, completes within minutes on a standard desktop computer. The different approaches yield a biologically meaningful grouping of the eight cancer types according to the estimated signatures (Fig. 1).

We have applied this approach to the characterization of kidney cancer and have shown that classification of subtypes according to mutational signatures is consistent with classification based on RNA expression profiling and mutation rates (Durinck *et al*., 2015).

## Acknowledgment

We thank Leonard Goldstein and Oleg Mayba at Genentech Inc. for their insights and suggestions.

## Funding

This work was supported by European Molecular Biology Laboratory, the NSF award “BIGDATA: Mid-Scale: DA: ESCE: Collaborative Research: Scalable Statistical Computing for Emerging Omics Data Streams”, and Genentech Inc.

